# Recurrent Dynamics of Rupture Transitions of Single Giant Vesicles at Solid Surfaces

**DOI:** 10.1101/2020.06.08.140178

**Authors:** V.N. Ngassam, W.-C. Su, D. L. Gettel, Y. Deng, Z. Yang, N. Wang-Tomic, V. P. Sharma, S. Purushothaman, A. N. Parikh

**Affiliations:** University of California, Davis; University of California Davis

## Abstract

Single giant vesicles (GVs) rupture spontaneously from their salt-laden suspension onto solid surfaces. At hydrophilic surfaces, they rupture via a recurrent burst-heal dynamics: during burst, single pores nucleate at the contact boundary of the adhering vesicles facilitating asymmetric spreading and producing a “heart” shaped membrane patch. During the healing phase, the competing pore closure produces a daughter vesicle. At hydrophobic surfaces, by contrast, the GVs rupture via a distinctly different, yet recurrent, bouncing ball rhythm: Rendered tense by the substrate interactions, GVs porate and spread monomolecular layer on the hydrophobic surface in a symmetric manner. Here too, the competition from pore closure produces a daughter vesicle, which re-engages with the substrate. In both cases, the pattern of burst-reseal events repeats multiple times splashing and spreading the vesicular fragments as bilayer patches at the solid surface in a pulsatory manner. These remarkable recurrent dynamics arise not because of the elastic properties of the solid surface but because the competition between membrane spreading and pore healing, prompted by the surface-energy dependent adhesion, determine the course of the topological transition.

**STATEMENT OF SIGNIFICANCE:** Giant lipid vesicles adhering to a solid surface experience strong mechanical stresses. The contacting membrane segment loses thermal fluctuations and accumulates mechanical tension, the equilibration of which can give rise to global shape changes, lipid phase separation, and traction forces. Beyond a threshold tension, vesicles porate, unravel, and spread. Here, we find that a competition from pore-healing can make rupture iterative, rather than a single all-or-nothing event. During burst, single pores expand, spreading a lipid bilayer on the hydrophilic surface and a monolayer on the hydrophobic one. During heal, pore-healing can produce daughter vesicles. This burst-reseal event reiterates “splashing” portions of single vesicles at the solid surface and “bouncing” the remainder as a secondary vesicle in multiple steps.

## INTRODUCTION

Topologically closed giant vesicles (GVs) (1–4), isolating femto-to picoliter quantities of aqueous core from the surrounding bulk, are the simplest cell-sized compartments(2) (5-50 μm in diameter). They are delimited by a barrier membrane(5) (4-6 nm thick), which is a self-assembled single lamellae of phospholipid bilayers(6) held together by non-covalent hydrophobic interactions(7). Acting as a highly deformable elastic sheet, the membrane presents a unique combination of elastic properties: low shear modulus (~10^−3^ N.m^−1^), due to the in-plane lateral fluidity(8); large volume compressibility (~10^9^ −10^10^ N.m^−2^) and large area expansion (10^2^-10^3^ mN.m^−1^) moduli, reflecting the cooperative strength of the hydrophobic effect(9); and relatively low bending rigidities (10^−19^ N.m)(10), a few times larger than the thermal energy, ~20 *k*_B_*T* (where *k*_B_*T* ≈ 4 × 10^−21^ J). As a consequence, the membranes of GVs resist large-scale thickness fluctuations, bend readily, but tolerate only a limited area expansion (~5%) before the cumulated tension fails the membrane and ruptures the vesicle.

In the vicinity of an attractive solid surface, giant vesicles experience strong mechanical stresses. It is now well-established that the adhesion of GVs at the solid-liquid interface(11, 12) suppresses membrane undulations, rendering membranes mechanically tense(13); elevates intravesicular hydrostatic pressure and generates traction forces(14); increases membrane permeability(15) and drives shape transformations(11); and induce lateral phase separation of membrane components, thus stabilizing membrane heterogeneity(16, 17). In a limiting case, when the adhesion energy overcome the elastic energy penalty aganist the deformations that occur at the solid-membrane contact line, GVs rupture at solid surfaces(11, 18). Here, the balance between the adhesion energy per unit area, *W*, and the surface tension, *σ* – given by Young-Dupre’ equilibrium relation *W* = *σ*(1 – cos*θ*), where *θ* represents the contact angle at the adhesion rim - stretches the membrane beyond the rupture tension (2-5 mN/m) (19) promoting the formation of a membrane pore through a thermally activated, stochastic nucleation process(20). Once a pore opens, the membrane tension can continue to relax via pore expansion allowing the solid-membrane contact line to spread further, thus facilitating an irreversible topological transition rupturing the GV at the solid surface and forming a well-defined quasi-two dimensional, single lipid bilayer patch terminated by free edges.

The biophysical mechanisms by which single giant unilamellar vesicles rupture at solid surfaces have been recently studied. Using time-lapse fluorescence microscopy, Hamai and co-workers(21–23) monitored the time-dependent progression of vesicle destabilization at solid surfaces and identified multiple different pathways. At hydrophilic surfaces, they found that an overwhelming proportion of vesicles (~92% cases, n=39) followed a single mode, which they dubbed asymmetric isolated rupture pathway. Here, the rupture proceeds through the nucleation of a pore near the rim of the adsorbed vesicle at the substrate surface. A subsequent expansion of the pore and an abrupt rupture-induced spreading of the membrane (10-20 ms) then resulted in a characteristic heart-shaped bilayer patch.

At hydrophobic surfaces, by contrast, GVs wet the aqueous interface by a distinctly different mechanism. A hydrophobic surface in water is characterized by a large surface energy (~ 40-50 mN/m). This high interfacial energy then provides the driving force for the adhesion of the GV at a hydrophobic surfaces. Because the gain in the adhesion energy per unit area (*W*), comparable to the surface energy, far exceeds the rupture tension (2-5 mN/m) of the vesicular membrane, a GV wetting a hydrophobic surface ruptures and spreads. The spreading of the lipids from the source GV releases the hydrophobic surface energy by producing a monomolecular lipid layer, which transforms the hydrophobic interface into a hydrophilic one(23–25). The kinetic pathways characterizing the rupture of GVs at hydrophobic surfaces are reported recently by Zan and co-workers(26) using time-resolved video microscopy. Their results support a mechanism in which a rare event leading to the disruption of the outer leaflet of the adhering GV initiates the transfer of lipids to the hydrophobic surface thus creating a precursor “hemi-fusion diaphragm” at the contact line. This initial loss of lipids from the outer leaflet alone produce a mismatch in molecular densities of the two leaflets, which then tenses the outer leaflet and lowers the energy barrier for pore formation near the contact line. The pore-edges facilitate lipid exchange between the leaflets and their eventual transfer to the surface ultimately transforming the GV into a monomolecular lipid layer at the hydrophobic surface.

Although different, the two mechanisms characterizing behaviors of GVs at hydrophilic and hydrophobic surfaces are both mediated by the formation of a membrane pore, which forms due to the mechanical tension produced by the adhesive interactions between single GVs and the solid surface. Considering this central role of pore formation in driving vesicle rupture raises a general question: *How does the competition from pore healing mediate the rupture process?* It is clear that the formation and growth of a pore helps mechanically stressed vesicles relieve membrane tension both by reducing effective membrane area and decreasing the vesicular volume, promoting spreading(27, 28). But the pore formation also creates a solvent-exposed free edges. The reorientation of the edge lipids into a hemi-micellar configuration driven by hydrophobic forces(29), then accrues mechanical tension at the edge (γ). As a consequence, the internal Laplace pressure (= 2*σ*/*R*), where *R* corresponds to the radius of the deformed GV, facilitates the volume loss from the vesicular compartment and promotes membrane healing by pore-closure(27, 30). Thus, a balance of the competition between these two effects, spreading and healing(31), must dictate the dynamics and final morphologies produced by the rupture of vesicles at solid surfaces.

Here, we identify a new, heretofore unappreciated, feature of surface-mediated rupture of giant vesicles. We find that an interplay between spreading and healing can give rise to non-trivial dynamics characterizing vesicle rupture at the solid-liquid interface. Specifically, we find that when spreading effects due to adhesive interactions do not dominate, the pore-nucleated GV rupture process is no longer an all-or-nothing event; but rather it follows a well-orchestrated, iterative sequence of steps in a surface energy-dependent manner. On hydrophilic surfaces, single GVs rupture by the asymmetric rupture pathway, but the transformation of the three-dimensional vesicle into a membrane patch does not conclude with a single rupture. Instead, the rupture process involves a series of repeated burst-heal events characterizing the topological transformation: During the burst regime, single pores nucleating at the contact boundary of the adhering deflated vesicles not only produces the “heart” shaped patches of lamellar membrane fragments but also gives rise free daughter vesicles likely by the healing of non-adhering portions of the parent GV. This burst-reseal event reiterates multiple times “spilling” portions of single GV membranes at the solid surface producing multiple heart-shaped patches of supported membrane patches. On low energy hydrophobic substrates, we find that the symmetric rupture, which occurs rarely on hydrophilic substrates, becomes the dominant pathway for vesicle rupture. Here too, we observe that the GV rupture is recurrent. It proceeds via a trampoline-like bouncing ball rhythm in which single vesicles striking the substrate form microscopic pores and spread partially at the point of contact between essentially undeformed GVs and the underlying hydrophobic substrate before detaching from the surface and bouncing-off. The process repeats multiple times before the vesicle membrane is transformed into a lamellar monolayer adhering firmly to the hydrophobic surface. In both cases, this cyclical pattern of poration, spreading, and resealing, arising from a competition between adhesion-mediated membrane spreading and pore healing, causes unusual dynamics characterized by wetting, splashing, and bouncing of giant vesicles at solid surfaces in a surface free energy - dependent manner.

## EXPERIMENTAL SECTION

### Materials

POPC(1-palmitoyl-2-oleoyl-*sn*-glycero-3-phosphocholine), Rhodamine B DOPE (1,2-dioleoyl-*sn*-glycero-3-phosphoethanolamine-N-(lissamine rhodamine B sulfonyl)) were purchased from Avanti Polar Lipids (Birmingham, AL, USA); Sucrose, Sodium Chloride, and Potassium Chloride were from Fisher Chemical (Fair Lawn, NJ, USA); Glucose, Toluene, Chloroform, Acetone, Sulfuric acid and Octadecyltrichlorosilane (OTS), 90+% were purchased from Sigma-Aldrich (Saint Louis, MO, USA); Hydrogen peroxide was from EMD Chemicals (Gibbstown, NJ, USA); Dulbecco Phosphate Buffered Saline 1X (DPBS) without Calcium chloride and without Magnum chloride was purchased from Gibco (Grand Island, NY, USA). All chemicals were used without further purification and all aqueous solutions were prepared with 18.2 mΩ-cm Milli-Q deionized water.

### Preparation of Giant Vesicles

Giant vesicles were prepared by adapting the well-established electroformation method developed by Angelova and co-workers(32). Small droplets (15–20 μ L) of lipid solution in chloroform (2 mg/ml) were spread on an ITO coated cover slip and allowed to dry under vacuum for at least 2 hours. The dried lipid cake was then hydrated with a 300 mM sucrose solution in deionized water and sandwiched using a second ITO slide. GVs were electroformed by subjecting the sandwich to A 4V AC sine-wave voltage at 10 Hz for 90 minutes followed by a 4V square wave voltage at 2 Hz for an additional 90 minutes.

### Preparation of hydrophobic substrates

Glass substrates (22 x 22 mm coverslips, Corning, Corning, NY) were cleaned for 3-5 min with piranha etch, a 4:1 mixture of sulfuric acid and hydrogen peroxide heated to 90°C, to remove organic residues. (Caution: this mixture reacts violently with organic materials and must be handled with extreme care.) The substrates were then rinsed copiously with deionized (18 mΩ-cm) water and dried under a stream of nitrogen. The deposition of *n*-octadecyltrichlorosilane (OTS, H_3_C(CH_2_)_17_17SiCl_3_) was achieved by adapting a previously described method(25). Briefly, all freshly oxidized, cover glass substrates were immersed in a 50 ml of 2.5 mM OTS solution in anhydrous toluene. The substrates were allowed to incubate in the solution for approximately 45 - 55 minutes. All silanization reactions were carried out in glass containers under nominally dry ambient conditions (relative humidity < 20%). After removal from the solution, the film-covered substrates were rinsed with chloroform and washed extensively with acetone under ultrasonic conditions to remove any excess reactants. Silanized samples were used within 1-3 days of preparation.

### Imaging of GVs incubated in different osmotic balanced concentrations of salty solutions

Aliquots (15μl) of freely suspended membranes of sugar-encapsulating GVs were studied at ambient temperature in either eight-well chambers fitted with a glass bottom coverslip (Nunc, Rochester, USA) or closed chambers fitted with OTS coated glass coverslips. We chose chambers over the conventional cover slip sandwiches because (1) they reduce the risk of GV deformation by mechanical pressure; (2) they afford large sample volumes, which reduces the possibility of inadvertent generation of significant osmotic imbalances due to solvent evaporation during extended experimental timescales (e.g., overnight); (3) they enable osmotic gradient generation in real-time; and (4) they allow solvent exchange and bilayer formation directly inside the chamber. GVs were incubated for 10 to 30 min in 300 mM glucose solution for control, then in (1) 150 mM NaCl, (2) 150 mM KCl, or (3) mixtures of 150 mM glucose and 75mM NaCl.

### Fluorescence Microscopy Measurements

Vesicles were monitored in real-time using either epifluorescence microscopy (Nikon Eclipse TE2000 inverted fluorescence microscope (Technical Instruments, Burlingame, CA) equipped with a Roper Cool Snap camera (Technical Instruments) and a Hg lamp as the light source) or a fluorescence microscope equipped with spinning disk confocal configuration using an Intelligent Imaging Innovations Marianas Digital Microscopy Workstation (3i Denver, CO) fitted with a CSU-X1 spinning disk head (Yokogawa Musashino, Tokyo, Japan) and a QuantEM512SC EMCCD camera (Photometrics Tuscon, AZ). Fluorescence micrographs were obtained using oil immersion objectives (Zeiss Fluor 40x (NA 1.3), Zeiss Plan-Fluor 63x (NA 1.4), and Zeiss Fluor 100X (NA 1.46); Carl Zeiss Oberkochen, Germany). Rhodamine-B DOPE (Ex/Em; 560/583) was exposed with a 5 mW 561 laser line. ImageJ (http://rsbweb.nih.gov/ij/), a public-domain software, and Slidebook digital microscopy imaging software (3i Denver, CO).

### FRAP Measurements

Fluorescence recovery after photobleaching (FRAP) experiments were performed with a 5 mW 561 laser line. Vesicles were viewed in the equatorial plane with a 40x objective and a QuantEM512SC EMCCD (electron-multiplying charge-coupled device) camera, giving a 512 x 512 pixel image. Rho-B DOPE fluorescent probes were bleached in a circular region (~3.2 μm in radius) at 100% of maximal laser power. Recovery of fluorescence was recorded and measured from the subsequent 250 frames, 10 frames per second. The diffusion coefficient was calculated as 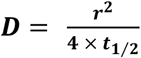, where r is the radius of the bleached spot and ***t*_1/2_** is the half-time of recovery(33).

## RESULTS AND DISCUSSION

### General Remarks

For our study, we used electroformed giant vesicles (GVs, 20-50 μm in diameter)(4) consisting primarily of a common unsaturated phospholipid, namely 1-palmitoyl 2-oleoyl-*sn*-1-glycero-3-phosphocholine (POPC) and encapsulating 100-300 mM sucrose under room temperature (~23.0 (± 0.5) °C) conditions (**Experimental section**). For real-time visualization of the membrane by time-resolved fluorescence microscopy, we doped the GVs with a small concentration (≤1.0 mol%) of a probe lipid, namely N- (lyssamine *Rhodamine* B sulfonyl)-1,2-dioleoyl-*sn*-3-phosphatidyl ethanol amine (Rho-DOPE) (34). Transferring the sucrose-laden GVs to an osmotically balanced bath containing isomolar concentrations of less dense glucose facilitates their gravitational settlement onto the underlying surface. Under these conditions, the GVs gravitating toward the underlying surface, hydrophilic and hydrophobic alike, remain largely undeformed retaining their spherical shape and exhibiting bending-dominated and thermally-excited topographical undulations (**Supporting Video 1**). By contrast, when the extra-vesicular dispersion medium, although osmotically balanced, contained even a small concentration of sodium chloride (≥ 5 mM), the GVs adhere to the underlying surface, likely by overcoming the hydration, undulation, and electrostatic repulsion(18). In what follows, we describe the behaviors of GVs at hydrophilic and hydrophobic surfaces as revealed by time-lapse fluorescence microscopy.

### Rupture of Single GVs at Extended hydrophilic Surfaces

A representative time-lapse video (**Supporting Video 2**) and a corresponding montage of selected frames (**Fig. 1a**) of epifluorescence micrographs reveal salient features of the dynamics characterizing the morphological destabilization and topological transitions of giant vesicles at hydrophilic (i.e., glass) surface. Consistent with previous reports(21, 22), we found that immediately upon contacting the surface (~10-20 s), the GVs undergo an abrupt topological transition over sub-second time scales abandoning their topologically closed morphology and producing characteristic heart-shaped patches adhering to the substrate surface. Surprisingly however, we found that this vesicle-to-patch transition is not a single, catastrophic event – a feature not appreciated previously(21, 22). Specifically, we found that a single daughter GV detached from the surface and diffused in the solution before re-engaging with the substrate surface merely tens of seconds later. In majority of the cases, the daughter GVs migrated several micrometers away from the initial patch before re-interacting with the bare substrate and producing a secondary heart-shaped patch. Interestingly, this process repeated several times before exhausting the vesicular material below the optical detection limit.

**Figure 1.**
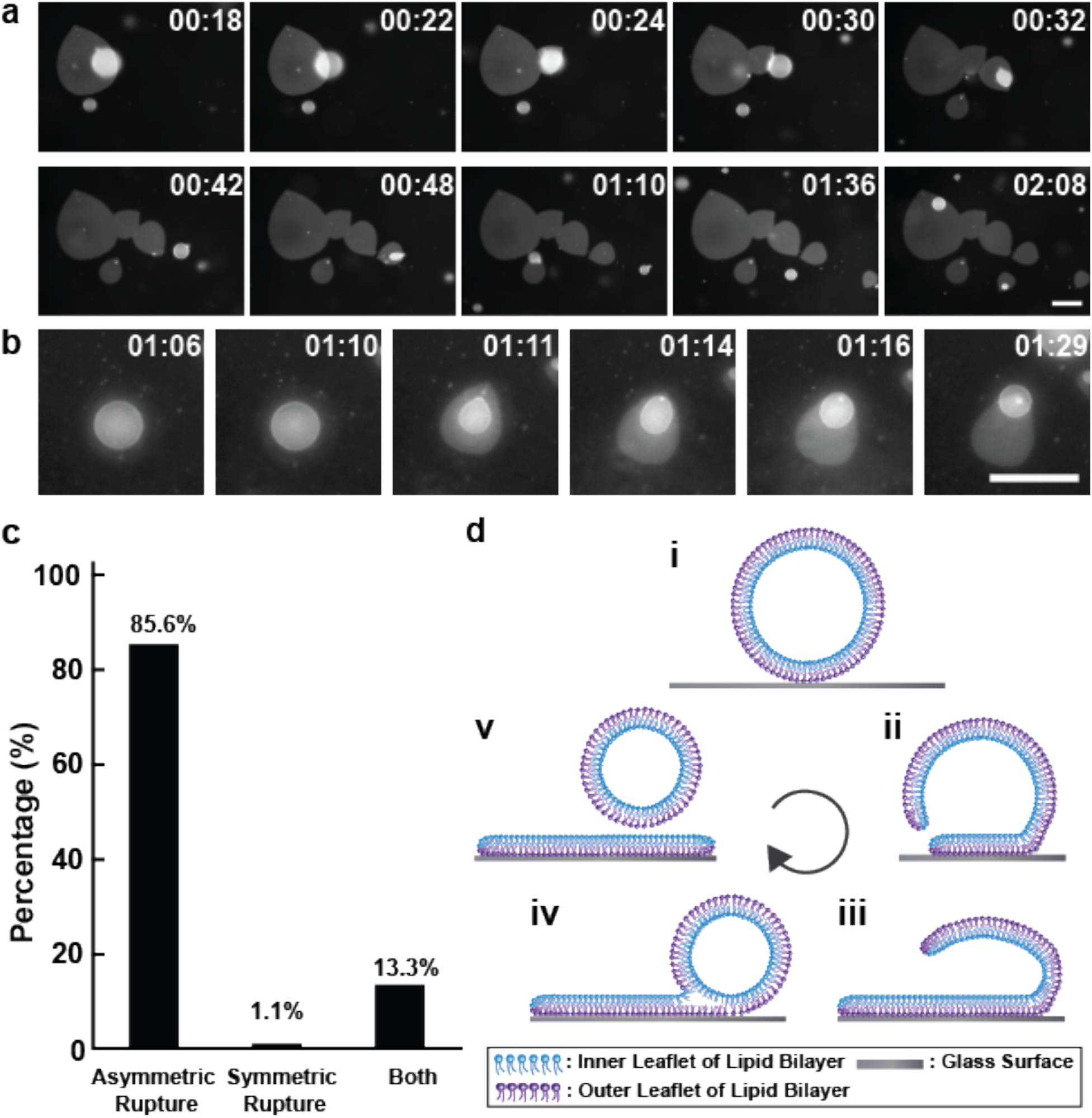
Dynamic rupture features of GVs settling onto high-energy hydrophilic glass surface. (a) Selected frames from a time-lapse video of fluorescence images of a single GV on glass substrate reveal multiple burst and reseal cycles. The GVs were 98% POPC doped with 2% Rho-DOPE. They contained 300mM sucrose in their interior and were immersed in an aqueous solution containing 150 mM glucose and 75 mM NaCl (b) Selected frames from a video of fluorescence images of a single GV on glass substrate showing a single burst and reseal cycle. (The GV composition and conditions were same as in (a) except for 150mM NaCl in the bath.) Time stamp shows minutes: seconds. Scale bar: 50 μm. (c) Histogram shows percentages of GV rupturing through asymmetric pathway, symmetric pathway or both. 459 vesicles consisting 98% POPC doped with 2% Rho-DOPE were analyzed. (d) A schematic representation of GV rupture on hydrophilic glass surface.

This iterative nature of vesicle-to-patch transition was highly reproducible. Examining 459 independent events in thirty-two (n=32) independent experiments, we found two major pathways (**Fig. 1a, 1b**). *First*, an overwhelming 85.6% GVs exhibited the repetitive rupture behavior described above (**Fig. 1c**). *Second*, a smaller, 13.3%, proportion of the rupture events revealed a conspicuous difference. The daughter vesicles did not migrate sufficiently far from the bilayer patches to encounter bare substrate surface (**Supporting Videos 3 and 4, Fig. 1b and Fig. S1**). In all of these cases, the daughter vesicles fused with the extant patch transforming the initial heart-shaped patches into more symmetric oblong- or pear-shaped lamellar membranes. Interestingly, the rupture of GVs here did not follow the abrupt asymmetric pathway. Rather, a gradual decrease in vesicle diameter and correspondingly a gradual expansion of the existing patch ensued, such as occurs on hydrophobic surfaces (see below). The surface-bound membrane patches resulting from the topological transition exhibited lateral fluidity, comparable to that of fluid supported bilayers(35), as revealed by the diffusional characteristics of probe lipids (0.8 ± 0.2 μm^2^/s) in microscopy-based fluorescence recovery after photobleaching (FRAP) measurements (36) (**Supporting Materials, Fig. S2 and Supporting Video 5**).

There are two distinct scenarios that can explain the recurrent rupture dynamics above: (**A**) successive ruptures of different membranes of structurally complex GVs and (**B**) multiple partial ruptures of single membrane leading to partial deposition of vesicular lipids at the substrate surface with the residual remodelling into a daughter vesicle. The former requires that the GVs are either multilamellar, consisting of concentrically arranged lamellae in a cylindrically smectic (onion-like) organization or oligovesicular, consisting of a hierarchy of internal “organelle” vesicles(37, 38). While the electroformation method used in the present study has been shown to yield predominantly unilamellar vesicles (>80-95%), a minority of GVs exhibiting internal tethers and oligovesicular organization, are invariably produced(39–41). Thus, the possibility that the recurrent rupture dynamics we witness arises from sequential ruptures of different lamellae or from the internal organelle vesicles merits consideration. Several independent lines of evidence support the (**B**) scenario. *First*, we note that although wide-field fluorescence images of GVs in **Fig. 1** do not permit us to directly discriminate between unilamellar and multilamellar vesicles, the large proportion of GVs exhibiting multiple ruptures (~86%) already suggests that GVs with complex architectures may not be required.

*Second*, to further consider the possible role of structural complexity of GVs in facilitating multiple ruptures, we used confocal fluorescence microscopy (**Supporting Video 6 and Fig. 2**). We examined three different GVs, each of which displayed no internal structure and presented a uniformly-lit, sharp fluorescence boundary consistent with unilamellar architecture. The montage of images shown in **Fig. 2** reveal an abrupt change in vesicle size at different times, which we interpret to correspond to the secondary vesicles produced after the first rupture event in each of the three GVs. Notably, these secondary vesicles are considerably smaller than the original GVs: Their perimeters estimated at roughly 40% (n=3) of those of the initial GVs (**Supporting Video 6 and Fig. 2**). This transition from the initial size to the final size is also accompanied by a transient shape change characterized by the short-lived appearance of non-spherical oblong shaped GV lasting several seconds. Although confocal imaging precludes simultaneous observation of the GVs and the patch forming at the underlying glass surface, the observed change in GV sizes (and the corresponding transient shape dynamics) are fully consistent with the scenario **B** where the rupture process induces only a partial loss of vesicular material with the remainder reassembling into secondary daughter vesicles. The alternate scenario **A** invoking complete “unbinding” of single layers from multilamellar “onion phase” vesicles should result in comparable sizes of patches, which was not observed. Moreover, because these GVs did not contain internal structures, we can also rule out that the multiple ruptures originate from successive depositions of different membranes of an oligomeric vesicle. (Note also that oligomeric vesicles should release all internal vesicles after the first rupture producing multiple patches in the second step, which was also not observed.). *Third*, consistent with the scenario **B**, we find a conservation of membrane surface area. Specifically, we find that *V_n_* ≅ *P*_*n*+1_ + *V*_*n*+1_ where *V_n_* corresponds to the surface area of the precursor GV (n^th^ step), *V*_*n*+1_ is the area of the secondary daughter vesicle produced in the *n*+1 step and *P*_*n*+1_ represents the area of the membrane patch (**Supporting Materials, Fig. S3**).

**Figure 2.**
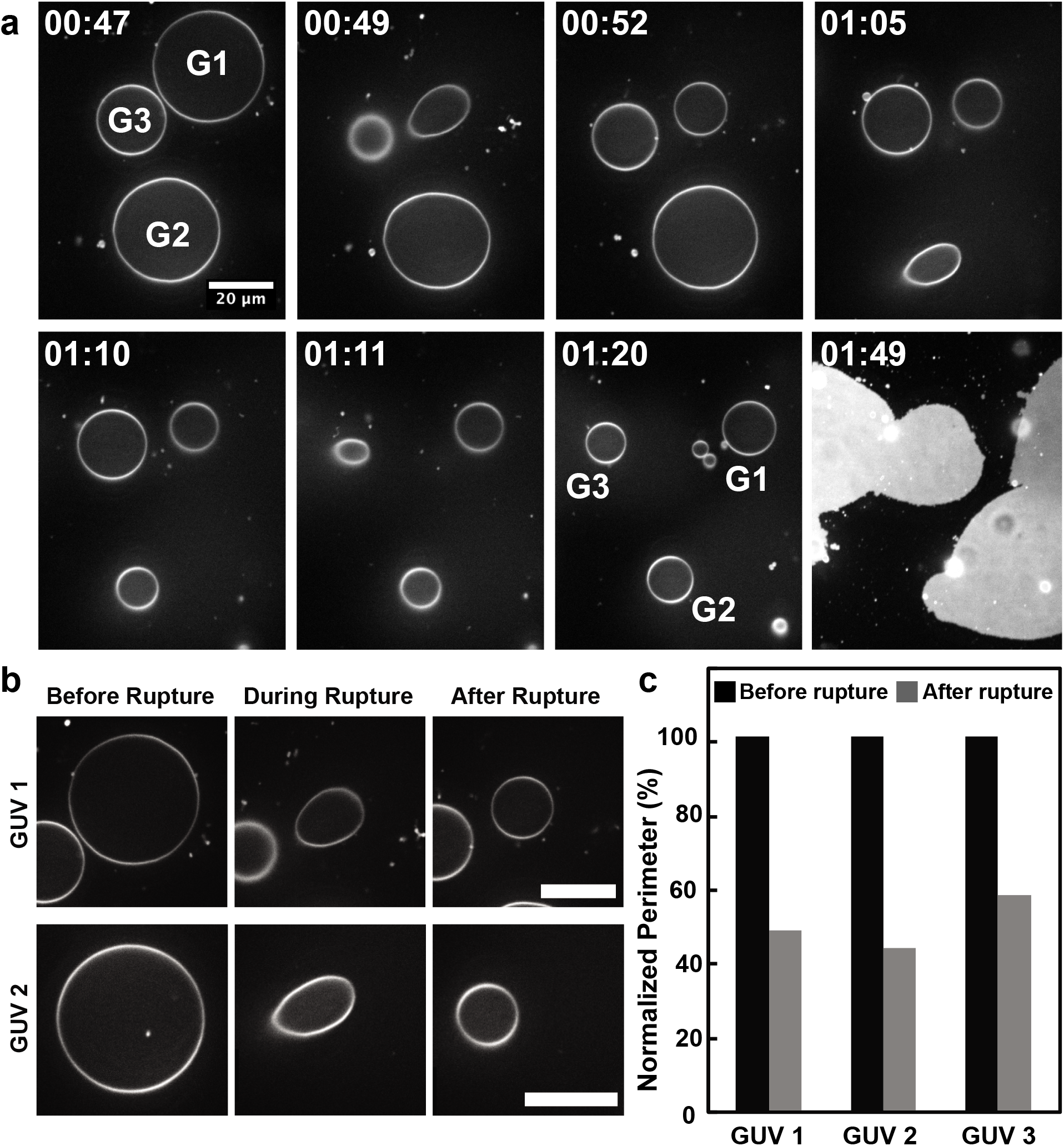
Significant size reduction of GVs after primary rupture event. (a) Selected frames from a video of confocal fluorescence images of GVs consisting of 98% POPC and doped with 2% Rho-DOPE containing 300mM Sucrose immersed in 150 mM NaCl solution incubated on glass substrate. Time stamp shows minutes: seconds. Scale bar: 20 μm. (b) Selected frames from (a) highlighting deformation and size reduction of GVs before, during and after rupture. (c) Histogram represents normalized perimeter of 3 individual vesicles at the equatorial plan before and after the first rupture.

Taken together, the results above can be reconciled in terms of a picture of vesicular rupture as a multi-step process. We propose that it arises from a competition between tendencies for membrane spreading(25, 42) and pore-healing triggered by adhesion-mediated membrane poration(30, 43). It is orchestrated by a sequence of biophysical events including substrate mediated adhesion, poration, spreading, and resealing - together generating the rupture dynamics such as described below.

At the aqueous interface of hydrophilic surfaces, the behavior of GV follows a well-defined course. It begins with the approach of the GV from the bulk solution to the glass surface. The presence of salt (> 100 mM) in the aqueous solution serves to lower the electrostatic repulsion (Debye-Hückel length, *κ* < 1 nm) between the negatively charged glass surface at the solution pH (~5.6) and the essentially zwitterionic GVs containing ~1 mol% negatively charged fluorescent probe(44). At the point of contact, the adhering GV deforms to produce a flat interface or a quasi-hemispherical cap maximizing its contact area (*A*) with the surface thereby gaining adhesion energy (*-WA*). The corresponding cost in curvature energy, determined by the bending rigidity, *k*, becomes manifest as a contact curvature(12), 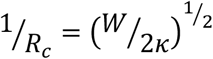, where *R_c_* is the contact curvature along the meridian of the GV. Taking *κ* = 10^−19^ J for POPC membranes and *W* = 10^−6^ – 10^−3^ J/m^2^ for moderate to strong adhesion(11), the radius of the contact curvature estimates at a few tens to a few hundreds of nanometers (14 ≪ *R_c_*(*nm*) ≤ 400). Because the free energy of membrane deformation is inversely proportional to the square of the radius of curvature, the contact curvature accrues significant curvature stresses locally near the rim of the interface between the GV and the solid surface(18). Moreover, the membranes of the adhering GVs experience an elevated membrane tension(20). These two properties, i.e., the global membrane tension and the local curvature stresses at the rim, collaborate to lower the activation energy barrier for the nucleation of a membrane pore(43). Although tension-mediated pore formation is a thermally activated, stochastic process(19, 45) and the pore may, in principle, form anywhere in the membrane, the presence of local curvature stresses preferentially localize the pore formation to the vicinity of the rim(31). Furthermore, to avoid the loss of adhesion energy, it seems reasonable that the pore nucleates and spreads preferentially in the non-adhering membrane side of the rim. The free edges of the open pore then expand under the influence of the internal Laplace pressure, a process further stabilized by the adhesion with the underlying solid. This in turn drives spreading away from the site of pore formation giving rise to the characteristic heart-shaped membrane patch. These considerations have been previously invoked to explain the spreading of GVs in single rupture events(21, 22).

This picture of adhesion-mediated membrane rupture however does not account for the repetitive rupture of single GVs giving rise to multiple membrane patches, such as we observe (**Fig. 1**). This requires that the adhesive patch separate and secondary vesiculation ensue. This scenario can be understood by considering the competition between pore expansion and pore healing(28, 30). Specifically, the pore formation creates solvent-exposed free edges. The reorientation of the edge lipids into a hemi-micellar configuration(29) driven by hydrophobic forces then generates mechanical tension at the edge (γ), which for a typical lipids is on the order of tens of piconewtons(46). This edge tension can then provide a driving force for membrane healing by pore-closure(27). In the present case, we propose that the pore healing can occur by the joining of the edge of the unfused membrane with the membrane at the boundary separating the fused and the unfused segments of the vesicular membrane (**Supporting Materials, Fig. S4**). Because the fused segments of the membrane are mechanically tense with reduced thermal fluctuation, a substantial tension gradient can be expected at the junction separating the two segments. We speculate that this tension gradient can provide the energy needed to surpass the energy barrier for the topological division required to divide a single contiguous membrane to produce a secondary, daughter vesicle(31) – mechanically akin to the process of tension-mediated membrane fission(47). The release of the daughter vesicle then allows for its diffusion into the bulk solution, and the subsequent approach to the surface. The process then repeat itself producing multiple heart-shaped membrane patches until curvature stresses and membrane tension are insufficient to induce poration. Whether GVs rupture in single or multiple events then depend on the balance between the two membrane spreading and secondary vesiculation processes, both of which depend sensitively on the membrane-surface adhesion energy.

### Rupture of Single GVs at Extended Hydrophobic Surfaces

Replacing the hydrophilic glass surface by a hydrophobic surface revealed a drastically different path by which GVs rupture at solid surfaces. A representative time-lapse video and a corresponding montage of selected frames of epifluorescence micrographs (**Supporting Video 7** and **Fig. 3**) of GVs settling onto OTS derivatized, hydrophobic glass surface (**See Materials and Methods**) documents this progression. Unlike the heart-shaped patches, which formed almost instantaneously when GVs settled onto hydrophilic silica substrates, a gradual and radially symmetric spreading of the membrane lasting as long as 400-500 s was evident. The areal spreading of the membrane estimated at 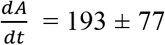 S.D. μm^2^/s, n=3, is of the same order of magnitude reported previously for the spreading of the more fluid, DOPC membrane (738 ± 29 S.D. μm^2^/s)(26). By contrast, the occasional and rare symmetric spreading on hydrophilic surface occurred at 2.6 μm^2^/s (**Supporting Materials, Fig. S5**) – roughly two orders of magnitude lower. This is not surprising because the spreading of lipids on a high-tension hydrophobic surface can be expected to release a considerably more interfacial free energy than at the hydrophilic surface(48) immersed in an aqueous bath (See below).

**Figure 3.**
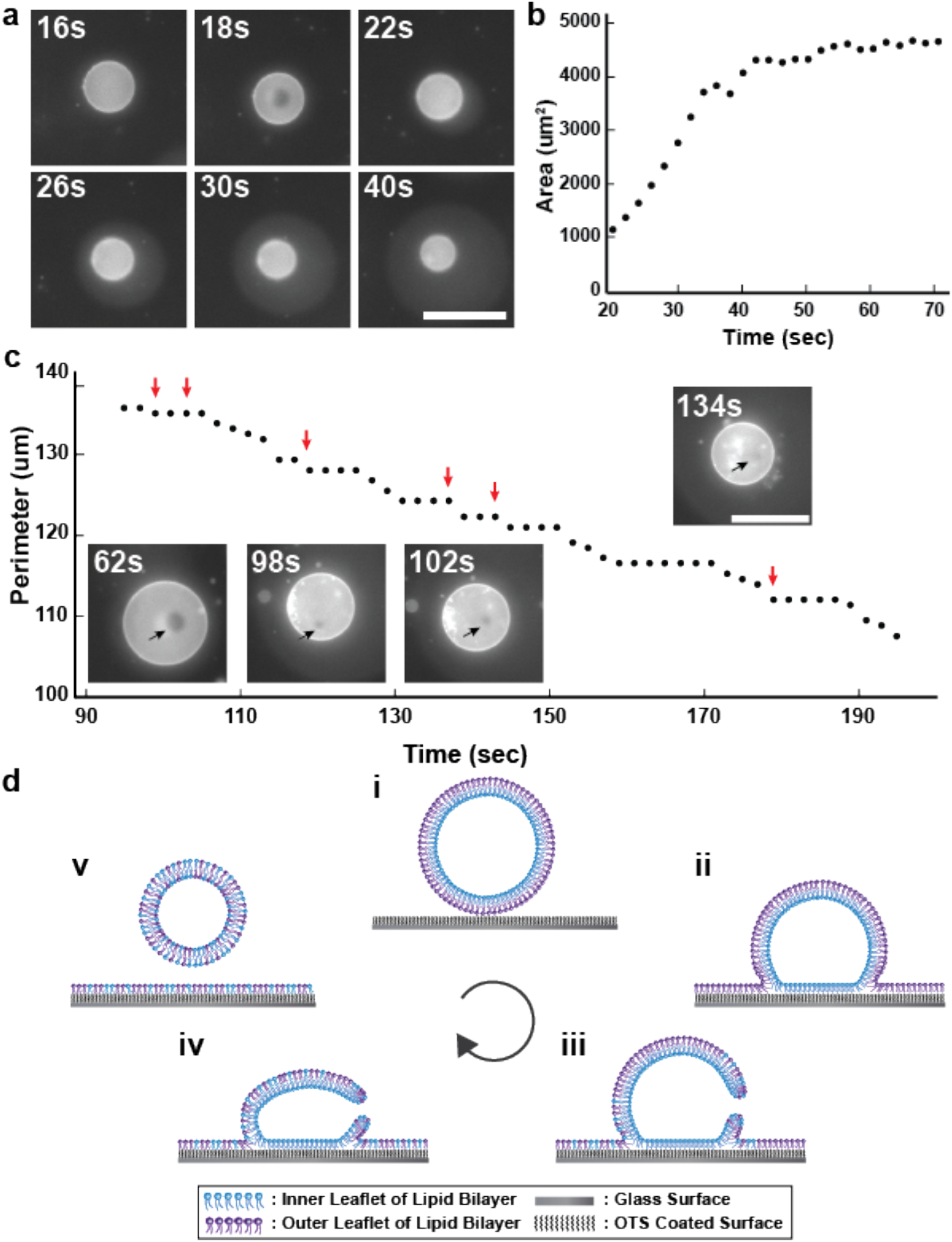
Dynamic rupture features of GVs settling onto hydrophobic surface. (a) Selected frames from a video of fluorescence images of a single GV consisting of 98% POPC and doped with 2% Rho-DOPE containing 300mM Sucrose immersed in 150 mM NaCl solution incubated on hydrophobic substrate. Scale bar: 50 μm. (b) Area measurements of symmetric spreading membrane area as a function of time. (c) Perimeter measurements of a vesicle while ruptured onto hydrophobic surface. Selected images show pore of a single vesicle ruptured on the substrate. Scale bar: 50 μm. Arrows are guides to the pore presented in video. (d) Schematic representation of GVs rupture on hydrophobic surface.

A closer view of the GV and the surface provided additional details of the vesicle rupture process. Examining the fate of the GVs during its interaction with the hydrophobic surface, we see that the parent GV largely retained its overall spherical shape (at the optical length scale) and exhibited a gradual decrease in their size. An analysis of the time-lapse images revealed the decay of vesicle size was not strictly monotonic (**Fig. 3c**), but rather that the vesicle size decreased in a step-wise manner creating a staircase-shaped profile. In each step, vesicle diameters decreased gradually for about 20-30 s before reaching the next plateau, which lasted for about 10-15 s. Moreover, at each plateau or each rung of the staircase profile, a single microscopic (2-4 μm in diameter) transient pore (**Supporting Video 8, Fig. 3c**), lasting hundreds of milliseconds, became visible. Comparing multiple steps of the single vesicles further revealed that the pores appeared randomly over the non-adhering part of the vesicle. Together, these observations indicate that vesicle rupture at hydrophobic surface is also not a single-step event. Instead a series of ruptures, each releasing a fraction of the membrane material from the parent GV to the solid surface, characterize vesicle rupture. In other words, the spreading of GVs onto a hydrophobic surface also followed a recurrent rupture dynamics: the membrane spreads, stalls, and spreads again.

The spreading of lipids at the high-energy aqueous interface of hydrophobic solids proceeds under a tension gradient (Δ*σ*) reminiscent of Marangoni flow(49). Previous studies have established that the balance between this Marangoni stress and the frictional dissipation at the spreading front then characterizes the dynamics of lipid spreading (Δ*σ* = *ζv*, where *ζ* is the frictional co-efficient and *v*, the spreading velocity)(42, 48). This yields an inverse square root dependence of spreading velocity on time: 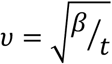, with a kinetic spreading coefficient 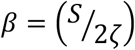 where *S* represents the spreading power given by the difference in free energy between lipids on the surface and those in the precursor GV. Adapting previously reported analyses of spreading kinetics, the time-dependent (1) radius of the circularly spreading monolayer on hydrophobic surface is 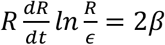 (49) and (2) area is 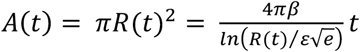, where *ε* represents the linear size of the initial contact zone at the onset of lipid spreading (Detailed Derivation in Supporting Materials, S2). Fitting the experimental *A*(*t*) v time data to the expressions above yield the values of the kinetic spreading co-efficient *β* between ~40 μm^2^/sec (*ε* = 0.5 *um*) and ~18 μm^2^/sec (*ε* = 5 *um*). (**Supporting Materials, Fig. S6**). These values of *β* shed light on the balance of forces. We begin by recalling that the OTS monolayer, which defines the hydrophobic surface is covalently attached to the underlying glass. Thus, the spreading of lipids at the OTS surface encounters frictional resistance at the methyl-methyl interface. Approximating the friction co-efficient *ζ*, between OTS and the lipid monolayer by that between the leaflets of lipid bilayers(50), which has been variously estimated to be between 1 x 10^8^ and 3 x 10^9^ Ns.m^−3^ (51–53), our estimates for *β* translate into a rather broad range of values for the spreading power, *S* between 4 and 8 mN.m^−1^ for *ζ* = 1 x 10^8^ Ns.m^−3^ and between 120 and 240 mN.m^−1^ for *ζ* = 3 x 10^9^ Ns.m^−3^. These values are clearly larger than the range of rupture tension values (2-6 mN.m^−1^) of equilibrated lipid bilayers, suggesting that the lipid spreading should be favored at the hydrophobic surfaces.

The considerations above suggest a mechanistic picture for the behavior of GV at hydrophobic surfaces, such as proposed below. A hydrophobic surface in water has a large surface energy(54). The interfacial energy between the OTS surface presenting a densely packed, two-dimensional lattice of −CH_3_ groups and water is 40-50 mN/m, which is considerably higher than expected from simple estimation of van der Waals forces(55). Since this adhesion energy is much higher than the rupture energy (5-8 mN/m)(19), thermodynamics favor the rupture of GVs and the coverage of the hydrophobic surface with a monomolecular layer of lipids. From a dynamic consideration, the GV rupture requires membrane poration. The structure of water in the close vicinity (*λ* ≈ 1 *nm*) of the high-tension, extended hydrophobic surface is known to be highly layered(54, 56), oriented with unsatisfied hydrogen bonds directed toward the surface(57), and one significantly reduced and fluctuating equilibrium density(7). Thus, it is reasonable that the GV approaching a hydrophobic surface experiences an abrupt hydration gradient *R* > *λ*. Since hydration also affects packing of lipids(58) in the vesicular membrane, a plausible consequence of the gradient in hydration across the vesicular dimension is the lowering of the activation energy required to nucleate a defect in the water-exposed outer leaftlet. This then initiates a sequence of molecular processes(26), which explain the topological transition in which GVs abandon their closed topology furnishing lipids needed to produce the monomolecular layer covering the hydrophobic surface in water. Specifically, the surface energy of the hydrophobic surface provides the driving force, i.e., the spreading power, allowing the spreading of the lipids from the outer leaflet. This must necessarily tense the outer leaflet. The mechanical tension in the outer monolayer together with the mismatch in molecular densities across the two leaflets fosters the formation of a membrane pore. The pore relaxes the membrane tension and allows for the two competing processes: (a) pore-mediated lipid exchange between the two monolayers and continued spreading under the influence of the spreading power and (b) pore healing promoting pore closure. These two processes then explain the recurrent spreading we observe.

#### GV Rupture at Mixed Wettability Patterned Surfaces

The drastically different processes by which GVs rupture, spread, and heal at hydrophilic and hydrophobic surfaces naturally raises an intriguing question: *How do GVs rupture at amphiphilic surfaces consisting both of hydrophilic and hydrophobic regions?* To address this question, we prepared surfaces presenting binary, microscopic spatial patterns of surface wettability consisting of alternating 100 μm (roughly 5 x the size of a representative GV) wide stripes of hydrophilic silica and hydrophobic surfaces. Monitoring the behavior of single GVs at these surfaces in real-time (**Fig. 4, Supporting Video 9**) revealed an unexpected behavior characterized by three salient features. *First*, the GV rupture became localized almost exclusively to the pattern boundaries separating hydrophilic and hydrophobic regions of the solid surface. The origins of this enhanced attractiveness for GV adhesion at the wetting boundary are at present not clear to us. We speculate the role of wettability gradient. A hydrophilic surface immersed in water is uniformly covered by water whereas the aqueous interface of a macroscopic hydrophobic surface is essentially “dry.” In other words, at the spatial boundaries of chemically patterned surfaces, water structure must undergo a rather drastic structural change. This in turn produces an epistructural tension(59) – the reversible work required to span the aqueous interface of the patterned surface – which provide a driving force for an enhancement in GV adhesion. *Second*, the vesicles bound to the pattern boundaries at the substrate surface, quite remarkably, spread in a strikingly asymmetric fashion preferentially covering the hydrophilic regions of the patterned surface. A consequence of this spatially patterned mode of vesicle-surface interaction is the formation of elliptically deformed bilayer patches on the hydrophilic channels, lithographically defined by the geometric properties of the underlying amphiphilic surface. We reason that this selection is kinetic arising from faster spreading kinetics at hydrophilic surfaces(25, 48). *Third*, in this mode of GV rupture and spreading, we find little or no evidence for the repetitive spreading indicating that the spreading overwhelms the competition from the pore healing (see above). Based on these characteristics, we propose that chemically structured surfaces or hydrophilic surfaces impregnated with hydrophobic “impurities” may foster GV adhesion, localize GV binding at pre-determined spatial locations, and suppress the recurrent rupture dynamics, thereby promoting complete membrane spreading in single bursts.

**Figure 4.**
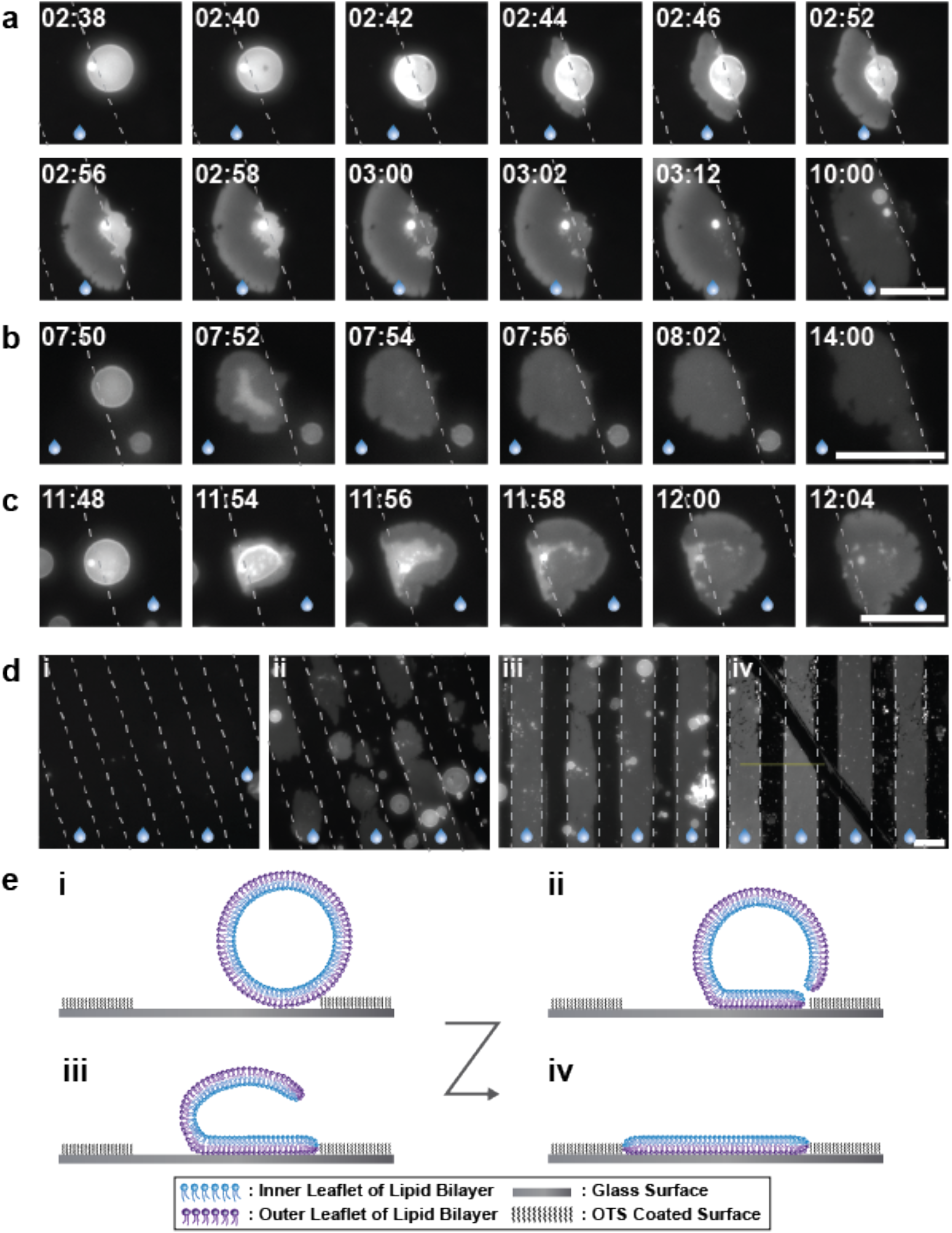
GVs destabilization on a pattern of hydrophilic and OTS coated hydrophobic substrate. (a-c) A time-lapse sequence of fluorescence images of different GVs consisting of 98% of POPC and doped with 2% Rho-DOPE containing 300 mM sucrose upon immersion in the external dispersion medium containing 150 mM NaCl solution and incubated on a pattern of hydrophilic and OTS coated hydrophobic substrate. Time stamp shows minutes: seconds. Scale bar: 50 μm (d) Image of the patterned surface (i) before, (ii) after 5 min, (iii) after 35 min of incubation of the GVs on pattern substrate and (iv) after rinsing with water on the patterned surface. Dash lines are guide to interface between hydrophilic and hydrophobic area. Water droplets are guide to hydrophilic area. Scale bar: 50 μm (e) Schematic representation of GVs rupture on the patterned hydrophilic and hydrophobic substrate.

## CONFLICTS OF INTEREST

Authors declare no conflicts of interest.

## AUTHOR CONTRIBUTIONS

All co-authors contributed to the design of the study. V. N. N. led the effort. V. N. N. trained the undergraduate researchers (YD, ZY, NW-T, VPS) and performed initial experiments. S. P. quantified the spreading rates. W. S. reproduced initial results, carried out additional control experiments, analyzed data, and co-wrote the manuscript.

## ACKNOWLEDGMENT

This work is supported by a grant from National Science Foundation DMR-1810540. The 3i Marianas spinning disk confocal used in this study was purchased using National Institutes of Health Shared Instrumentation Grant 1S10RR024543-01. We thank the MCB Light Microscopy Imaging Facility, which is a UC-Davis Campus Core Research Facility, for the use of this microscope. We are grateful to P. Rangamani for her careful and insightful critique of the manuscript.

